# Versatile NTP recognition and domain fusions expand the functional repertoire of the ParB-CTPase fold beyond chromosome segregation

**DOI:** 10.1101/2025.10.10.680097

**Authors:** Jovana Kaljević, Kirill V. Sukhoverkov, Katie Johnson, Antoine Hocher, Tung B. K. Le

## Abstract

Nucleotide triphosphate (NTP)-dependent molecular switches regulate essential cellular processes by cycling between active and inactive states through nucleotide binding and hydrolysis. These mechanisms were long thought to rely exclusively on ATPase or GTPase proteins, until the discovery of CTPase activity in the bacterial chromosome segregation protein ParB. In the ParAB*S* system, CTP binding enables ParB accumulation around the centromere-like *parS* DNA sites to activate the ATPase ParA, thereby facilitating chromosome partitioning to daughter cells. CTP hydrolysis then releases ParB from DNA for recycling. This discovery uncovered a new regulatory principle, but the broader diversity of proteins employing a CTPase mechanism remains unclear. Here, we conduct a large-scale survey of proteins harboring the ParB-CTPase fold across bacteria, archaea, bacteriophages, and eukaryotes. While many ParB-like proteins follow the canonical ParAB*S* organization with ParA partners, we also identify numerous orphan homologs encoded outside of the *parAB* operon, frequently linked to mobile genetic elements that may have driven their rapid diversification. The ParB-CTPase folds in these divergent proteins are often fused to lineage-specific domains with diverse predicted biological activities. We further demonstrate that while many homologs retain CTP-binding, others instead bind ATP or GTP, revealing a broader spectrum of nucleotide specificities than previously appreciated. Our findings establish the ParB-CTPase fold as a widely distributed and evolutionarily versatile NTP-binding module, repeatedly co-opted through domain fusion and shifts in nucleotide specificity to enable functions far beyond the classical ParAB*S*-mediated DNA segregation.

## Introduction

Molecular switches play central roles in biology, enabling cellular processes to be switched on and off at the right place and time inside cells. The switching mechanism is commonly mediated by nucleotide binding and release/hydrolysis, which toggle proteins between active and inactive states with distinct biological activities. This well-established paradigm was thought, for decades, to rely exclusively on ATPases or GTPases^1–4^ to regulate processes ranging from cell growth and differentiation to protein synthesis, folding, transport, and DNA replication, segregation, and repair. This view was, however, challenged by the recent discovery of proteins that instead use CTP binding and hydrolysis to drive conformational switching, revealing an alternative regulatory principle in biology^5–7^. This newly discovered mechanism raises an important question: how widespread are CTP-dependent molecular switches, and in what biological contexts do they operate?

The best-characterized CTP-dependent switch is in the bacterial DNA segregation protein ParB, a core component of the ParAB*S* system^8–12^. This tripartite system consists of an ATPase ParA, a DNA-binding CTPase ParB, and DNA *parS* sites^13^ (Fig. 1a). ParB acts as a clamp whose state is controlled by CTP binding and hydrolysis. Upon CTP binding, ParB closes around DNA and spreads away from *parS* sequence (active state, Fig. 1b, i-ii)^5,12,14–17^. The repeated loading and spreading of ParB-CTP enable many clamps to accumulate around *parS*, creating a platform for interaction with the ATPase ParA and thereby driving DNA segregation^7,10,12,18–25^. CTP hydrolysis eventually re-opens the clamp and releases ParB from DNA (inactive state, Fig. 1b, iii)^16,17,26^. This on-off cycle recycles ParB, preventing it from becoming irreversibly bound on DNA or becoming distant from *parS*^16,17,19,26^. In this way, a high local concentration of ParB partitioning complexes is maintained, which is essential for stimulating ParA’s ATPase activity. Although variable in sequence, canonical ParB proteins share a conserved domain organization (Fig. 1c)^5,6,12,16^. The C-terminal domain mediates self-dimerization and, in some species, contributes to nonspecific DNA-binding^27–29^. This C-terminal domain is connected via a flexible linker to the central helix-turn-helix (HTH) DNA-binding domain that specifically recognizes *parS*^12,30^. The highly conserved N-terminal domain^18,28,31^ contains a ParA-binding motif (LG-R/K-GL), crucial for interaction with ParA and stimulation of its ATPase activity^22,32^. The N-terminal domain also harbors the ParB-CTPase fold, which includes three essential sequence elements: Box I (or C motif), Box II, and Box III (or P motifs), which altogether enable CTP binding and hydrolysis (Fig. 1d)^5,6,16–18,26,31,33,34^.

**Fig. 1.**
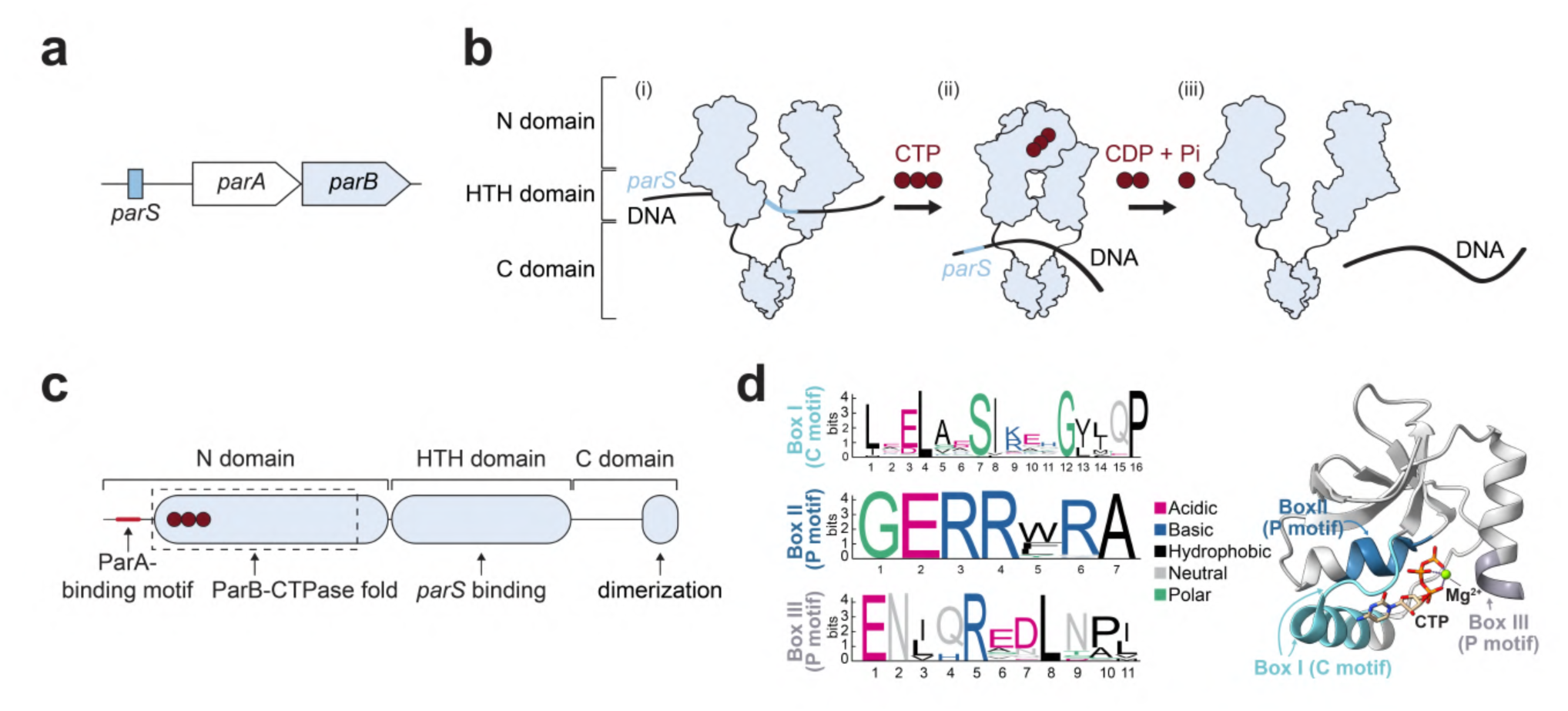
Canonical domain architecture, and CTPase mechanism of ParB proteins. **a** Schematic of the *parAB* operon with the upstream *pars* binding site. **b** Simplified model for ParB-CTP switch. (i) ParB binds CTP in an “open clamp” configuration and recognizes the *pars* sequence; (ii) **a** conformational change causes the clamp to close around the DNA, when ParB detaches from *pars* and spreads along the DNA; (iii) ParB eventually hydrolyses CTP to CDP and organic phosphate, which opens a clamp, leading to ParB’s release from the DNA. Domains are indicated on the left. c The N-terminal domain contains the ParA-binding motif (small red box) and the ParB-CTPase fold (dashed box, bound CTP shown as red circles). The central helix-turn-helix (HTH) domain mediates *pars* recognition, and the C-terminal domain mediates dimerisation. d Conserved sequence motifs and structure of the ParB-CTPase fold. Left: sequence logos for Box I (C motif, cyan), Box II (P motif, blue) and Box Ill (P motif, purple) derived from ∼1800 chromosomal ParB homologs; the height of the stack indicates the sequence conservation, while the height of symbols within the stack indicates the relative frequency of each amino acid residue at that position; amino acids are colored based on their chemical properties. Right: structural model of the ParB-CTP fold (PDB: 7BM8) bound to CTP and Mg2•, with conserved CTP-binding motifs (Boxes I, II and Ill) highlighted in cyan, blue, and purple, respectively.

Beyond canonical ParB proteins that are involved in DNA segregation, many homologs are also CTPases but function in other biological processes^12^. These proteins retain a similar domain architecture to canonical ParBs with an N-terminal ParB-CTPase fold, a central HTH DNA-binding domain, and a C-terminal dimerization domain. Some homologs also have a ParA-binding motif, such as PadC in *Myxococcus xanthus,* which binds CTP to recruit ParA to the bactofilin cytoskeleton and enhance the robustness of chromosome segregation^6,35^, and KorB from the multi-drug resistance RK2 plasmid in *Escherichia coli,* which instead uses CTP binding and hydrolysis to regulate long-range regulation of plasmid gene expression^36–38^. By contrast, other homologs retain the canonical domain organization but often lack the ParA-binding motif. For example, *Bacillus subtilis* nucleoid occlusion protein Noc uses CTP binding and hydrolysis to regulate not only DNA clamping/sliding but also its membrane-binding activity, coordinating chromosome segregation with cell division^30,39–42^. Other homologs, such as VirB from *Shigella flexneri*^12,43–48^, and BisD from *Pseudomonas putida* integrative conjugative elements^49^, employ CTP binding and hydrolysis to regulate gene expression. Despite their diverse roles, all these proteins act as CTP-dependent DNA clamps that load and slide along DNA, akin to the canonical ParB in DNA segregation^12^. Other ParB homologs, however, diverge significantly in both domain organization and function. For instance, the archaeal serine kinase SerK^50,51^, bacterial serine-kinase homolog SbnI^52^ ,and the eukaryotic sulfiredoxin Srx protein^53^ contain the ParB-CTPase fold but lack the HTH and C-terminal dimerization domains. Rather than acting as a CTP-dependent DNA clamp, their ParB-CTPase fold instead functions as an ATP-binding module essential for their enzymatic activities in serine biosynthesis and antioxidant metabolism, respectively^51,52,54^. These sporadic examples hint at the versatility of the ParB-CTPase fold. Yet, the full diversity, evolutionary distribution, and nucleotide specificity of this fold have not been systematically explored to date.

Here, we conducted a large-scale survey of proteins with a ParB-CTPase fold across bacteria, archaea, phages, and eukaryotes, uncovering a previously unrecognized breadth of domain architectures, including many novel domain combinations with clear lineage-specific patterns. By integrating biochemical assays with structural prediction, we show that while some variants retain CTP binding, others preferentially bind ATP or GTP, revealing a much broader spectrum of nucleotide specificity than previously recognized. Altogether, our findings expand the ParB-CTPase fold beyond the few characterized examples, demonstrating that this fold is a far more widespread and versatile NTP-binding module with potential roles in numerous biological processes.

## Results

### Orphan ParB proteins occur widely across diverse lineages

As ParB is the founding example of a CTP-dependent switch, we first surveyed the distribution of *parB* genes across prokaryotes, providing a baseline for identifying CTPases beyond the classical ParAB*S* segregation system (Fig. 2a). Using a balanced dataset of 64,060 bacterial and 4,649 archaeal genomes spanning 217 phyla^55^, we identified 80,326 ParB homologs (Fig. 2a and Methods). For each, we recorded whether *parB* was encoded on chromosomes, plasmids, or prophages, as predicted by Genomad. Finally, we took syntenic context into account to distinguish the canonical *parAB* operon involved in chromosome segregation from stand-alone *parB* genes (Figure 2a). We found that 84% of bacterial orders contained chromosomally encoded *parB* (Fig. 2a, blue bars in the inner ring, Supplementary Table 1), ∼5.6% of all *parB* homologs (4,542/80,326) were plasmid-encoded, and ∼1% (872/80,326) were prophage-encoded. We find that ∼77% (59,669/77,443) of chromosomally encoded *parB* genes are part of *parAB* operons in bacteria (Figure 2a, blue bars in the outer ring) (80% when we restrict our analysis to near-complete genomes). In contrast, while we find 4.8% of archaeal *parB* are found in the vicinity of a *parA* homolog, only 9 genomes satisfy our near-complete genome criteria, and none of these are complete. ParAB detection in archaea, therefore, likely reflects contamination, consistent with the general absence of chromosomal ParAB*S* systems in this domain of life, although plasmid-encoded *parB* homologs have been described in some archaeal lineages^56^.

**Figure 2.**
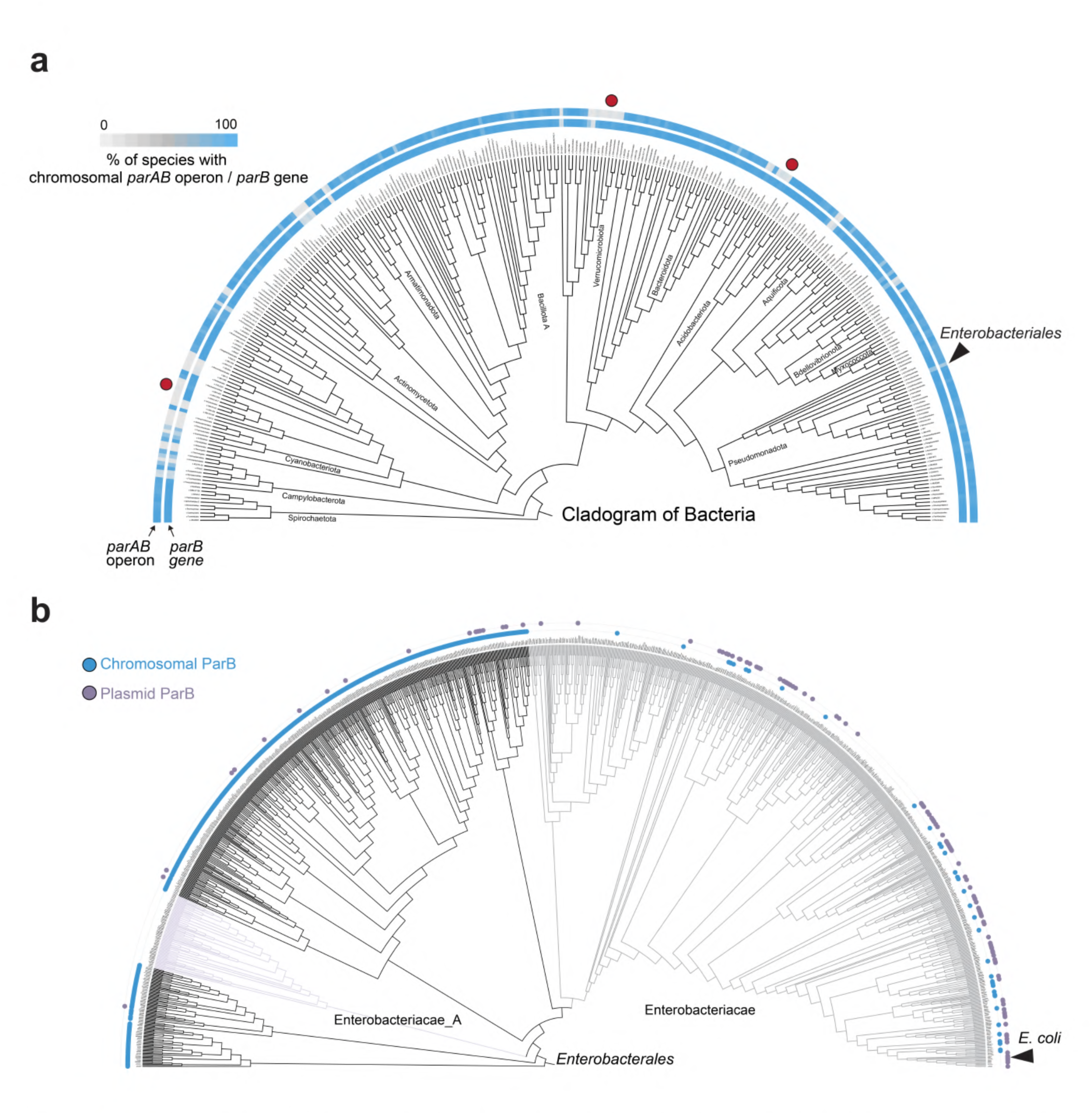
Distribution of the ParABS system across bacteria. **a** Frequency and distribution of chromosomal *parAa* genes across bacteria, summarized at the order level. Analyses were restricted to near-complete genomes (checkm2>95) and to orders for which we have a minimum of 7 near-complete genomes from different species (n=30842). The outer ring indicates whether a conserved *parAa* operon is present on the chromosome (blue=parAa present, grey=parAa absent). The inner ring marks species that encode a *para* gene, regardless of its genomic context (blue = *para* present, grey= *para* absent). Species with grey bars in the outer ring but blue bars in the inner ring represent orphan ParB proteins (i.e., where *para* is encoded without a *parA* partner), and selected cases are marked with red dots. Major bacterial clades are annotated. **b** Cladogram of species within the order *Enterobacteriales.* Each branch represents one species. Outer rings indicate the genomic location of the *para* gene: blue dots in the inner ring = chromosomally encoded *para;* purple dots in the outer ring = plasmid-encoded *para;* no dots in the inner and outer ring = no *para* detected. *E. coli* exemplifies a species that has lost chromosomal *para* but encodes the *para* gene on a plasmid, and its position is marked with an arrowhead.

While chromosomally encoded *parB* was found in most bacterial orders (Fig. 2a, blue bars in the inner ring), our survey uncovered numerous independent instances of *parB* loss (Fig. 2a, grey bars in the inner ring), which typically coincided with loss of the entire *parAB* operon (Fig. 2a, grey bars in the outer ring). Such losses occurred sporadically across phylogenetically and ecologically diverse bacterial clades, consistent with earlier findings that the ParAB*S* system is widespread but not universal^13^. For example, within *Enterobacteriales*, some species, including *E. coli*, retain *parB* on plasmids^8,57–61^ (Fig. 2b, black arrowhead and purple dots in the outer ring), whereas others lack it entirely from both chromosomes and plasmids (Fig. 2b, no purple and blue dots in the outer ring). Interestingly, in several lineages, the *parAB* operon has been lost, yet a ParB*-*like protein is encoded elsewhere on the genome (Fig. 2a, grey bars in the outer ring and blue bars in the inner rings, red dots). We term these orphan ParB proteins: homologs that retain clear sequence similarity to canonical ParBs, including the ParB-CTPase fold, but occur outside the conserved *parAB* operon. To position orphan ParBs within broader diversity, we constructed a single-gene tree for all bacterial and archaeal ParB and ParB-like proteins, annotating each for the presence or absence of ParA-binding motif (Supplementary Fig. 1). Across all sequences, the ParA-binding motif was strongly enriched in the subset of ParB proteins encoding in the vicinity of a *parA* homolog (hypergeometric test, p-value <1e-16), supporting their co-evolution. In fact, ParB proteins containing a ParA-binding motif cluster together (Supplementary Fig. 1, red dots in the outer ring), whereas those lacking such a motif scattered across bacteria and archaea (Supplementary Fig. 1, no red dots in the outer ring). These orphan ParB proteins include well-characterized examples such as Noc (Supplementary Fig. 1, magenta), which arose through gene duplication and neo- functionalization from a canonical *parB*^30,40^. Overall, our phylogenetic analysis indicates that orphan ParB proteins are both common and taxonomically widespread.

### A domain-centric search reveals the broader diversity of the ParB-CTPase fold

To explore how the ParB-CTPase fold extends beyond the canonical ParB proteins that function in DNA segregation, we conducted a domain-centric search using the ParB N-terminal domain (PFAM ID: PF02195) that contains the ParB-CTPase fold as a query. This search revealed two main groups of ParB-CTPase folds that mirror the full-length ParB phylogeny (Fig. 3a vs. Supplementary Fig. 1). The first group corresponds mainly to bacterial ParB homologs with the canonical domain organization (Fig. 3a, blue bars in the inner ring, red and orange dots in the outer rings). In these, the evolution of the ParB-CTPase fold is rather constrained, as reflected by their shorter branches on the domain centered tree. Within this group there are well-studied examples such as chromosomal ParB from *B. subtilis*^5,62^ (Fig. 3a, example 1), as well as Noc, which lacks the ParA-binding motif but overall retains the canonical domain architecture (Fig. 3a, example 2)^40^. The second group consists of long-branched, divergent clades that span bacteria, archaea, phages, and eukaryotes (Fig. 3a, no red dots in the outer ring). Among bacterial proteins in this group, genomic context analysis revealed their frequent occurrence on plasmids and prophages (Fig. 3a, black and purple dots in the outer ring). While some retain domain organization similar to canonical ParB in DNA segregation (Fig. 3a, example 3), they are almost always orphan variants lacking the ParA-binding motif (Fig. 3a, examples 3-4, no red dots in the outer ring). Their enrichment on mobile genetic elements suggests horizontal transfer might have shaped their distribution. Within this divergent branch, we also identified the ATP-binding proteins SerK^50^ and Srx^53^ (Fig. 3a, examples 5-6). Their branching alongside many uncharacterized orphan ParB proteins suggests that this group might contain additional ATP-binding and/or other atypical ParB-like proteins.

**Figure 3.**
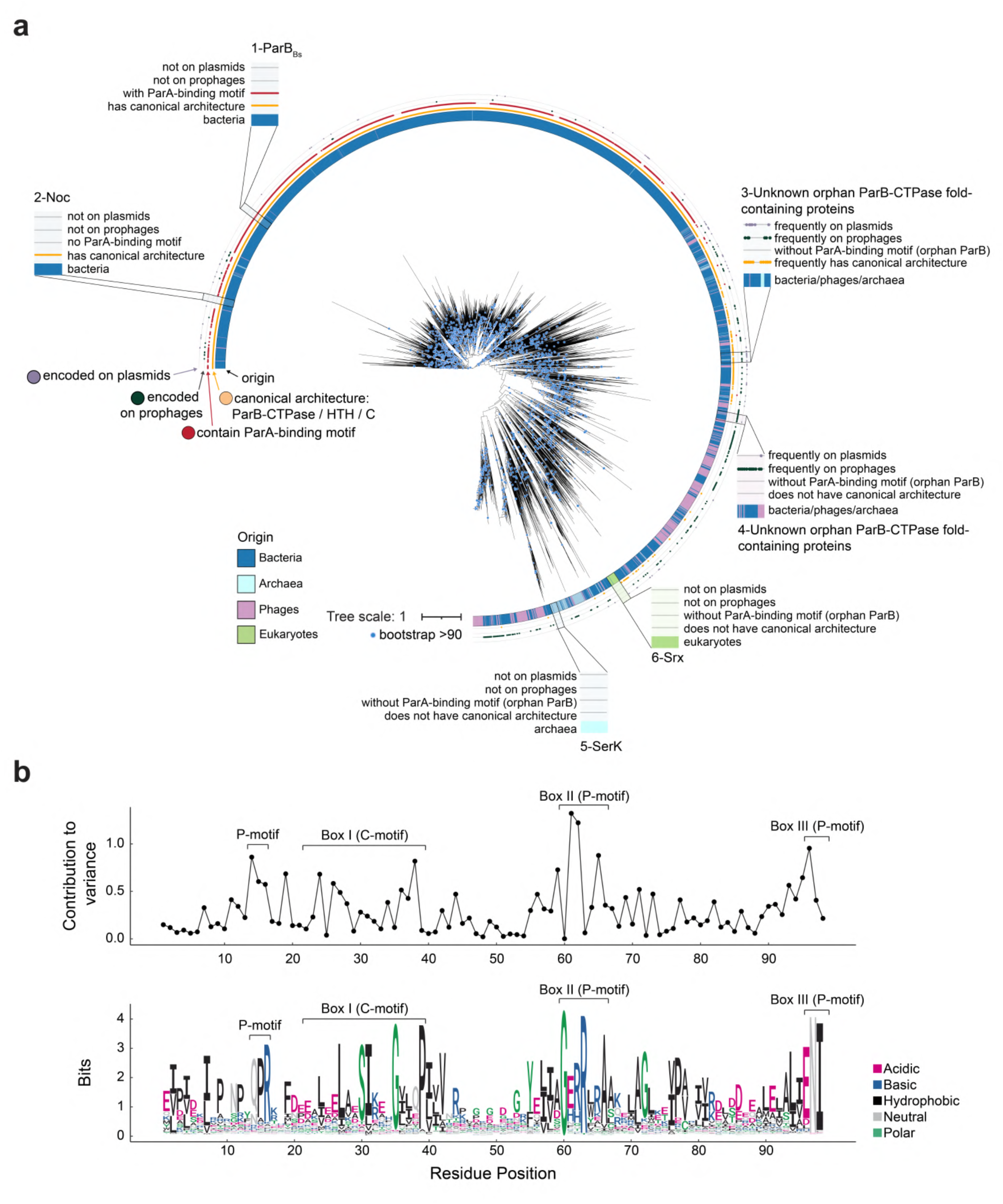
Phylogenetic diversity of ParB-CTPase fold-containing proteins. **a** Protein domain-level, maximum likelihood phylogenetic tree of 9,456 ParB-like domains from bacteria, archaea, phages, and eukaryotes (midpoint rooted); Bootstrap support 1:90 is indicated by blue dots. Concentric rings summarize key features (from inside to outside): the inner ring shows taxonomic origin (blue = bacteria, cyan = archaea, pink = phages, green = eukaryotes); the orange ring indicates canonical domain organization (defined as positive hit against TIGR00180); the red ring indicates presence of ParA-binding motif (defined as (LIFll)G[KIR]G(LIFII); two outer rings indicate the genomic context (purple dots= plasmid encoded genes; dark-grey dots = prophage encoded genes). Known examples are annotated for easier following, and they include: canonical ParB (1), Noc (2), SerK (5), and Srx (6). Novel orphan ParB-like candidates are highlighted in examples 3 and 4. b Sequence analysis of the ParB-CTPase domain. Top: per-residue contribution to sequence variability across the alignment (see methods), with conserved motifs indicated: Box I (C motif), Box II (P motif), Box Ill (P motif). Bottom: sequence logo of the same alignment showing residue conservation at each position; amino acids are colored based on their chemical properties.

Next, to pinpoint residues responsible for the divergence of the ParB-CTPase fold, we quantified the contribution of each amino acid to overall sequence variation and found that divergence is not distributed evenly across the ParB-CTPase fold but concentrated in discrete clusters (Fig. 3b). The strongest signals mapped to the C motif (Box I) and P motifs (Boxes II and III), crucial for CTP binding in canonical ParB^5,6,16,17^, raising the possibility that natural variations in these motifs might have altered ligand specificity, potentially driving the evolutionary diversification of ParB-like proteins. Altogether, our analysis so far reveals that the ParB-CTPase fold extends far beyond known examples, encompassing widespread yet largely uncharacterized classes of NTP-binding proteins.

### ParB-CTPase fold frequently associates with lineage-specific domains to form fusion proteins

Next, we examined the domain organization of ParB-like proteins and listed all PFAM-A domains associated with proteins containing the ParB-CTPase fold. We found that the ParB-CTPase fold is rarely encoded alone; instead, it is often fused to additional domains to form larger multi-domain proteins (Fig. 4a). In bacteria, the majority (∼70%) of proteins with this fold are fused to additional domains, whereas this proportion is lower in archaea (∼50%) and phages (∼30%) (Fig. 4a). When we looked at the remaining proteins annotated as “single-domain protein” (Supplementary Fig. 2a), we found clear differences across lineages in their length (Supplementary Fig. 2b). In bacteria, most of these proteins are long (Supplementary Fig. 2b, blue, ≥300 amino acids), much larger than the CTPase fold itself (∼100 amino acids), indicating that they likely contain additional domains absent from PFAM-A. By contrast, in phages and a subset of archaeal sequences, we observed very short proteins (∼100 amino acids) that consist almost entirely of the ParB-CTPase fold (Supplementary Fig. 2b, cyan and magenta arrowheads). These likely represent genuine single-domain proteins, hinting that the fold itself can act as an independent functional unit, like sulfiredoxin.

**Figure 4.**
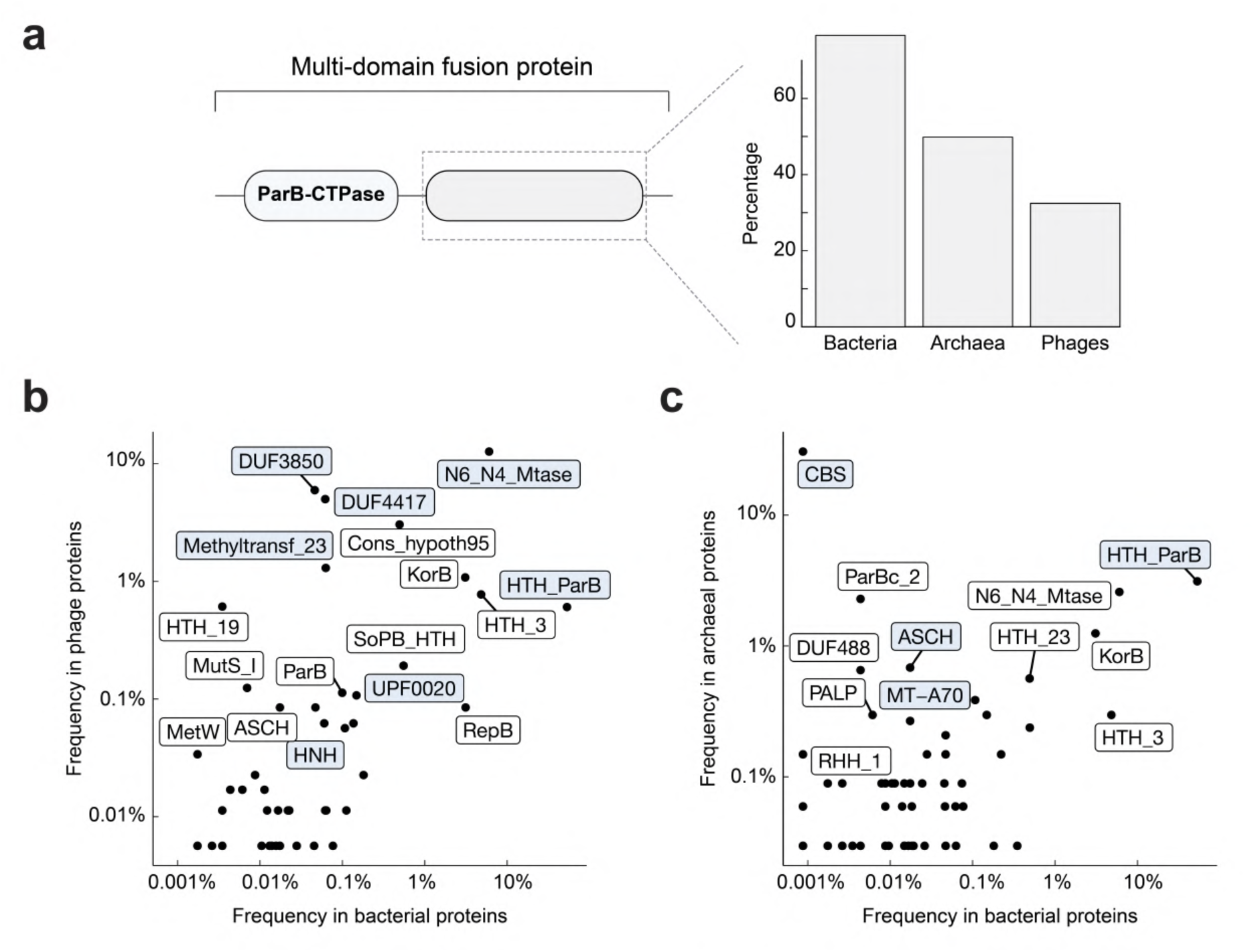
Domain organization of ParB-CTPase fold-containing proteins. **a** Many proteins containing the ParB-CTPase fold (PF02195) occur as multi-domain fusion proteins. Left: schematic of a multi-domain fusion protein showing ParB-CTPase fold combined with other domains. Right: proportion of ParB-CTPase proteins that are part of multi-domain fusions in bacteria, archaea, and phages. **b-c** Frequency of domains fused to the ParB-CTPase fold in bacterial (x-axis) compared to phages (panel b, y-axis) or archaea (panel c, y-axis). Each dot represents a distinct PFAM domain fused to the ParB-CTPase fold. Domains discussed in the main text are highlighted in light blue. **b** Domains that are more frequently found in bacteria are: HTH (PF17762), C-terminal dimerization domain (PF23552), HNH endonuclease (PF01844), and UPF0020-like (PF01170). Domains that are more frequently found in phages are: DUF3850 (PF12961), DUF4417 (PF14386), N6_N4 methyltransferase (PF01555), and Methyltransferase-23 (PF13489). c Same analysis for bacteria vs archaea. CBS (PF03471) is strongly enriched in archaea, as are ASCH (PF04266) and MT-A70 (PF05063), whereas HTH (PF17762) occurs in both domains of life. PFAM entry codes are provided for reference; See also Supplementary Fig. 3-4.

We next surveyed domains that are fused to the ParB-CTPase fold across bacteria, phages, and archaea, and found a clear lineage-specific pattern (Fig. 4b-c). In bacteria, ParB-CTPase folds are most often found in canonical ParB homologs, *i.e.,* fused to DNA-binding helix-turn-helix (HTH) and C-terminal dimerization domains (Fig. 4b-c, HTH). These fusion proteins were almost exclusively encoded on plasmids and chromosomes, consistent with their established role in DNA segregation^12,19^ (Supplementary Fig. 3a, grey arrowheads). We also discovered novel fusions, including ParB-CTPase fused to HNH domains (usually confer nuclease activity^63–65)^ or UPF0020 (a poorly understood family with predicted RNA-methyltransferase activity) (Fig. 4b), suggesting that bacteria harbor a broader functional diversity of ParB-like proteins than previously appreciated.

In phages, we found that ParB-CTPase folds were frequently fused with domains of unknown function (e.g., DUF3850, DUF4417) or domains with predicted DNA methyltransferase activities (e.g., N6N4-Mtase, Methyltransf23) (Fig. 4b). These fusion proteins were strongly enriched in prophage regions of bacterial genomes^66–69^, suggesting an important role in phage biology, perhaps in anti-defense or phage-host interactions^70–73^ (Supplementary Fig. 3a, dark green arrowheads). Strikingly, among all detected combinations of domain fusion, only the ParB-CTPase-DUF3850 fusion was consistently co-encoded with a neighboring *parA* homolog (Supplementary Fig. 3b). Since DUF3850 belongs to the PUA-like superfamily, typically associated with RNA recognition^68^, this unusual operon structure might suggest a previously unrecognized connection between ParA ATPases and RNA-related functions in prophages.

In archaea, the most frequent domain fusion was to the cystathionine-β-synthase (CBS) domain (Fig. 4c). CBS domains are known to regulate the activity of associated enzymatic or transporter domains in response to adenosyl-containing ligands such as AMP, ATP, or S-adenosylmethionine (SAM)^50,74–77^. Their fusion to ParB-CTPase fold suggests a possible coupling between nucleotide binding/hydrolysis and cellular metabolic states. The ParB-CTPase-CBS architecture was rare in bacteria, and within archaea, restricted mainly to Methanobacteriota and Haloarchaea (Supplementary Fig. 3c, purple dots in the outer ring), indicating vertical inheritance rather than horizontal transfer, consistent with its presence on chromosomes rather than mobile genetic elements. Archaea also encoded proteins with the ParB-CTPase fold fused to RNA-related domains such as the ASCH domain (RNA binding^78^) or MT-A70 domain (mRNA adenine methyltransferase^79,80^). In addition, we found canonical fusions with DNA-binding (HTH) and C-terminal dimerization domains, although these cases were scattered across archaea, and were rarely found in association with ParA (Supplementary Fig. 3c, orange dots in the outer ring), suggesting they are unlikely to function in chromosome segregation.

Next, to investigate whether ParB-CTPase co-varies with the different partner fusion domains, we compared sequence conservation across protein families (Supplementary Fig. 4a-c). Both the C motif (Box I) and P motif (in Box II) displayed subtle but systematic variation depending on the partner fusion domain (Supplementary Fig. 4a-c), suggesting that the nucleotide-binding pocket may co-evolve with the partner domain, potentially adjusting nucleotide specificity to perform specific biological functions.

### ParB-CTPase fold is capable of binding diverse NTPs, not limited to CTP, across domains of life

To examine nucleotide-binding properties of ParB-CTPase fold-containing proteins, we selected 50 diverse representatives spanning bacteria, phages, and archaea, prioritizing a broad range of domain architectures (Fig. 5a and Supplementary Fig. 5). These included fusions to domains of unknown function (e.g., DUF3850, DUF4417, and DU1015), DNA-binding (e.g., HTH), or enzymatic domains (e.g., acetyltransferases, SAM-dependent methyltransferases, and N6N4-methyltransferases). Of these, 28 proteins could be expressed and purified to homogeneity in high yield from *E. coli*. Eleven displayed NTP-binding activity in DRaCALA assays using radiolabeled ATP, CTP, GTP, and UTP. As controls, nucleotide-binding activity was abolished when the conserved arginine residue in the P motif (Box-II) was substituted by alanine (R→A mutants^5,6,16^; Fig. 5c-e, bottom rows), confirming that nucleotide recognition is mediated by the ParB-CTPase fold itself. Under our conditions, 17 candidates showed no binding (Supplementary Fig. 5). The lack of *in vitro* nucleotide binding in these proteins does not necessarily rule out ligand interaction *in vivo*, where binding may depend on additional cofactors such as DNA or protein partners.

**Figure 5.**
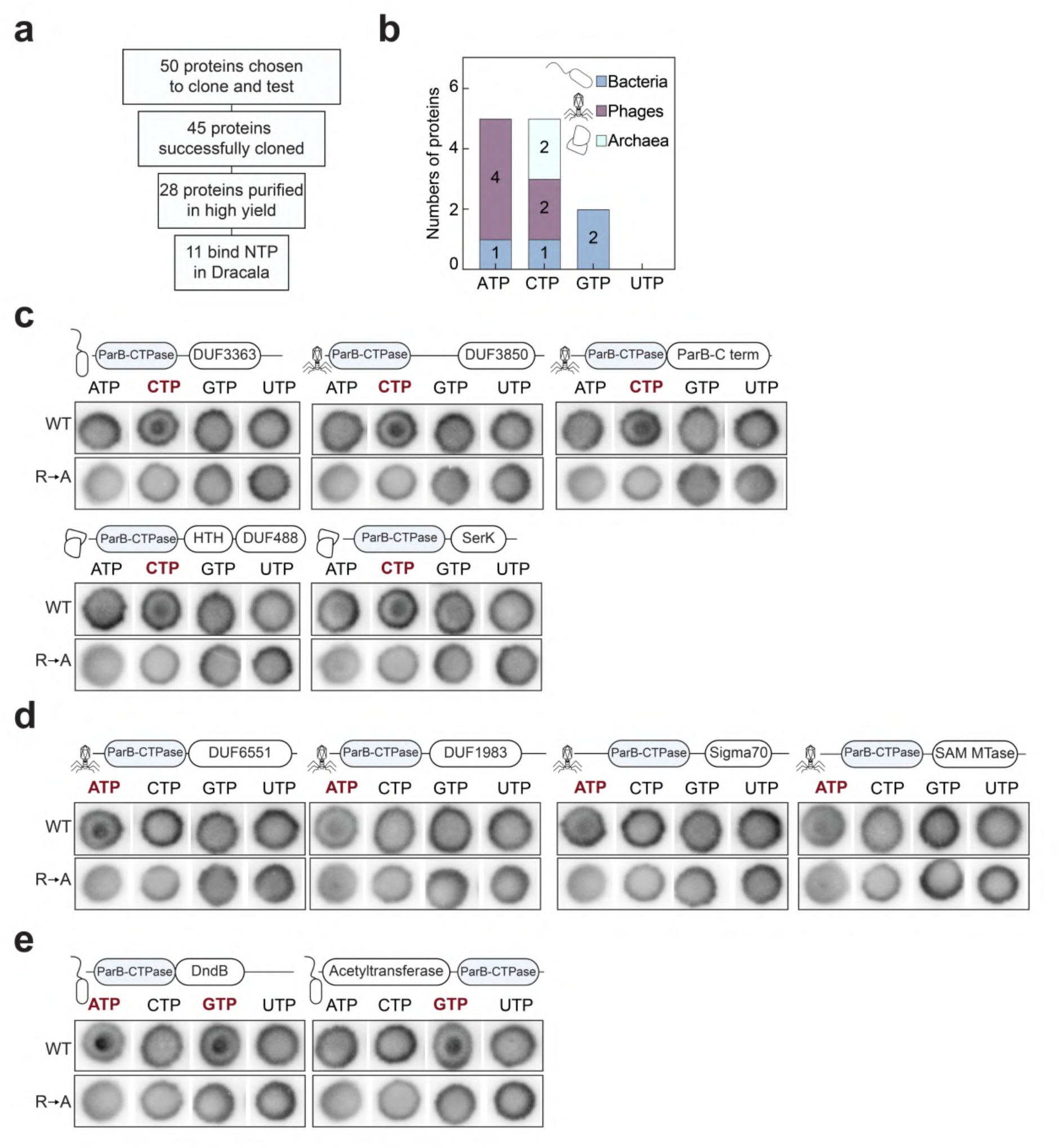
Screening ParB-CTPase fold-containing proteins for nucleotide binding using DRaCALA. **a** Experimental pipeline: of 50 proteins selected for testing, 45 were successfully cloned, 28 were purified in high yield, and 11 showed detectable NTP binding. **b** Summary of binding preferences; number of bacterial (blue), phage (magenta), and archaeal (cyan) proteins that bound ATP, CTP, GTP, or UTP in DRaCALA assays. c-e Wild-type proteins (WT, top) and R-->A mutants in the conserved P-motif (GxxR, negative control, bottom) were tested for binding to radiolabeled NTP o-P32. The bull’s eye staining indicates NTP binding due to a more rapid immobilization of protein-ligand complexes compared to free ligands alone. The starting concentration of proteins used was 25 µM. DRaCALA assays were performed with three replicates for WT and two replicates for R-->A mutants; one representative replicate is shown. Cartoon icons indicate the origin of each protein (phage, bacterium, or archaea). Domain schematics above each protein indicate representative fusions, with the ParB-CTPase domain shown in blue and fused to various domains. Proteins are grouped by their detected binding ligand: (c) CTP-binding proteins; (d) ATP-binding proteins; (e) GTP-binding proteins. In each panel, the binding ligand is highlighted in red above the DRaCALA blots. Detailed information for each protein (including source organism, accession numbers, and domain annotations) is provided in Supplementary Table 4. See also Supplementary Fig. 5.

Among the positive binders, we detected binding to CTP, ATP, and GTP (Fig. 5b). CTP binding was observed in five proteins (Fig. 5b-c): a DUF3363 fusion in bacteria; DUF3850, DUF6551, and a C-terminal dimerization domain fusions in phages; and a DUF488-HTH and SerK-like domain fusions in archaea. Notably, the SerK-like protein bound CTP, rather than ATP, in contrast to earlier reports^50^. ATP binding was detected in five proteins (Fig. 5b, d), including ParB-CTPase fold fusions to bacterial SAM-dependent methyltransferases (enzymes that transfer methyl groups to DNA, RNA, or other substrates using S-adenosylmethionine as a cofactor^81^ ) and Sigma70 domains (transcriptional regulation). In phages, ParB-CTPase fold fusions to two domains of unknown function bind ATP: DUF6551 and DUF1983, the latter frequently found at the C-terminus of phage receptor-binding proteins. Dual ATP/GTP binding was observed in a bacterial DndB-like protein (Fig. 5b, e). DndB is a ParB-like protein previously described as an ATP-dependent transcriptional repressor of the *dnd* operon, where through phosphorothiolation of DNA, it is involved in discrimination of foreign genetic elements^68,82–86^. Its ability to also bind GTP points to a possible additional, unexplored, layer in DndB biology. Finally, the only GTP-specific binder was a bacterial fusion of ParB-CTPase fold to an acetyltransferase domain (Fig. 5b, e), providing the first evidence of GTP recognition by a ParB-CTPase fold. Given that acetyltransferases catalyze the transfer of an acetyl group in numerous processes ranging from bacterial metabolism to chromatin regulation, nucleotide binding may modulate enzymatic activity or substrate specificity of these fusion proteins^87–89^. Altogether, our results demonstrate that the ParB-CTPase fold exhibits diverse nucleotide preference, leading us to wonder whether nucleotide specificity of the ParB-like fold could be predicted.

### Neither sequence analysis nor AlphaFold3 currently provide reliable predictions of nucleotide specificity

In canonical ParB proteins, nucleotide binding requires the coordination of three motifs: the C motif (Box I), which consists of one helix and one loop that contacts the nucleotide base, and P motifs (Box II and III), which contact phosphate groups and the magnesium ion (Fig. 1d). In non-canonical, orphan ParB proteins, Box III is often absent or highly divergent; thus, we focused our analysis exclusively on the C motif in Box I and P motif in Box II. Comparing sequences across CTP-, ATP-, and GTP binders showed that CTP and ATP binders have similar helix and loop length of the C motif (Supplementary Fig. 6a-c), with only minor variation (Fig. 6a, Supplementary Fig. 6a-c). The singular GTP-specific binder notably has a longer C motif helix (17 residues compared to ∼12 in ATP/CTP-binders, Fig. 6a and Supplementary Fig. 6a-c), although its loop length at the C motif was comparable with the ATP and CTP binders (∼6 residues, Fig. 6a and Supplementary Fig. 6a-c). The P motif in Box II was nearly invariable, spanning 6-7 residues across all binders (Fig. 6a and Supplementary Fig. 6a-c). Analysis of amino acid composition at C and P motifs revealed weak trends: C motif loops in CTP binders often contain mixed polar and hydrophobic residues (e.g., GLLQS, GLQEP, QQFFP, Supplementary Table 6), while those in ATP binders more frequently included bulky or hydrophobic residues such as phenylalanine, valine, or leucine (e.g., KDVPP, EFFLPTS, GFFKP, Supplementary Table 6) (Fig. 6a and Supplementary Fig. 6a-c). This trend was subtly reflected in hydrophobicity counts, with ATP-binding loops averaging ∼2.6 hydrophobic residues compared to ∼1.8 in CTP binders (Fig. 6b). Nonetheless, the modest biases could not reliably separate CTP, ATP, and GTP binders, indicating that simplistic sequence inspection does not easily enable prediction of nucleotide specificity.

**Figure 6.**
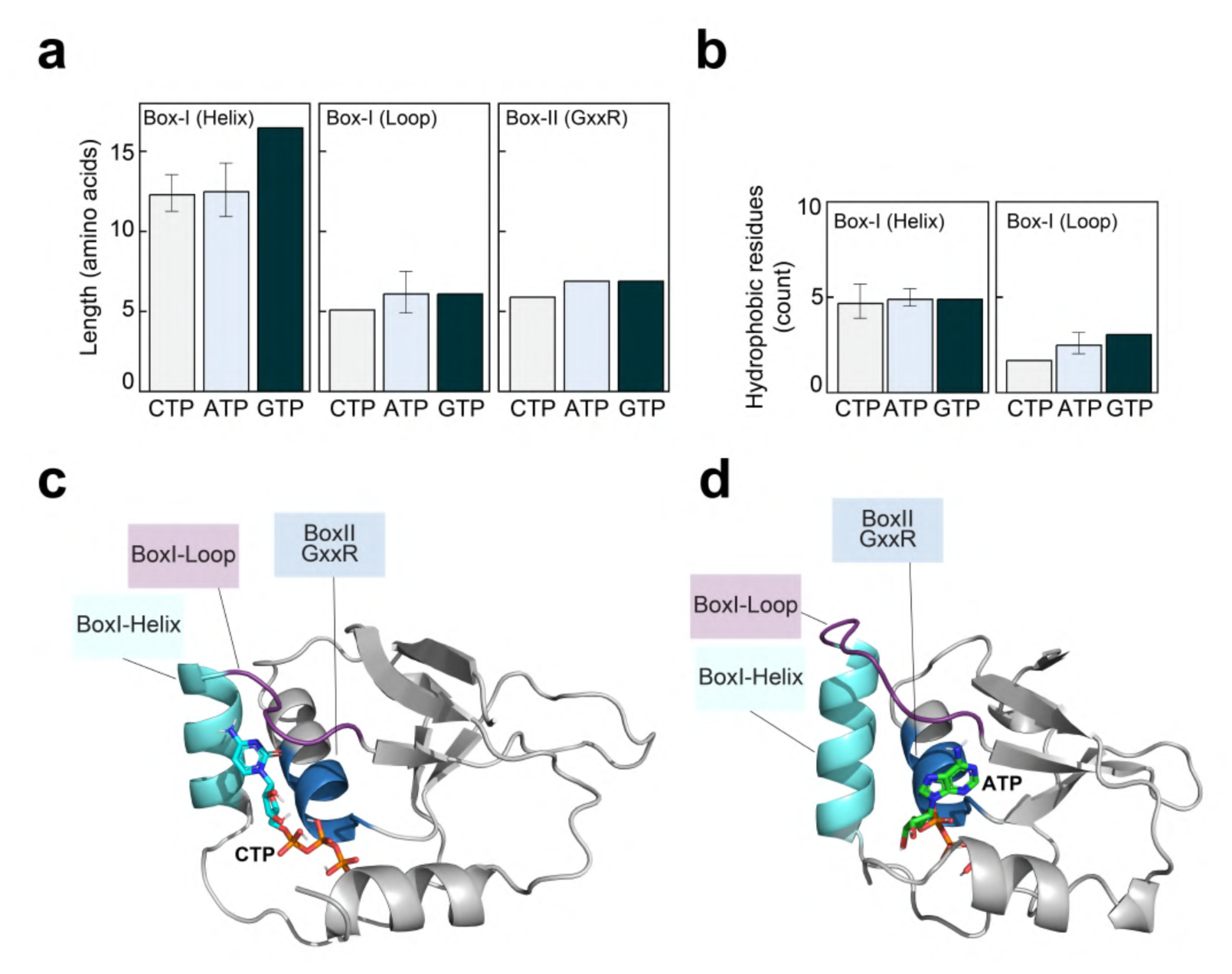
Sequence and structural features of the ParB-CTPase fold. **a** Average length (amino acids) of Box I (C-motif) helix, Box I (C-motif) loop, and Box II (P-motif, GxxR) region in CTP-, ATP-, and GTP-binding proteins. Bars show mean± standard deviation (sd). **b** Average number of hydrophobic residues in the Box I (C motif) helix and Box I (C motif) loop across CTP-, ATP-, and GTP- binding proteins. c Representative AlphaFold-predicted structure of a CTP-binding ParB-CTPase protein, showing the Box I (C motif) helix (cyan), Box I (C motif) loop (purple), and Box II (P motif, GxxR) motif (blue). Bound CTP (orange sticks) positioned towards both the helix and the loop. **d** Equivalent structure for an ATP-binding ParB-CTPase protein, showing the same motifs with bound ATP. ATP is positioned towards the loop only, and it seems not to contact the Box I (C motif) helix. See also Supplementary Fig. 6-8.

We then asked whether *in silico* protein-ligand co-folding using an in-house installation of AlphaFold3^90^ could enable the prediction of nucleotide preference. AlphaFold3 models of the above 11 positive binders with all four NTPs, followed by molecular docking, consistently placed nucleotides in the correct ligand-binding pocket (Fig. 6c-d, Supplementary Fig. 7). However, docking energies failed to distinguish between nucleotides, often ranking non-cognate ligands as equal or even more favorable than cognate ones (Supplementary Fig. 8a-c). Thus, AlphaFold3-based modeling currently cannot reliably reproduce the binding preferences observed *in vitro*. Still, closer analysis of AlphaFold3 models in complex with the cognate ligands revealed subtle trends (Fig. 6c-d). In CTP binders, cytosine base was frequently contacted by both the C motif helix and loop (Fig. 6c and Supplementary Fig. 7). In ATP binders, by contrast, the C motif helix is rarely observed to contact the base; instead, adenine was mainly contacted by residues from the C motif loop (Fig. 6d and Supplementary Fig. 7a). These patterns, however, were not universal, as the adenine base in several ATP binders (e.g., DndB-like and SAM-methyltransferase fusions) also contacts the C motif helix. Their C motif helix and loop sequence resembled those of CTP binders (Supplementary Fig. 7b). The sole GTP-specific binder showed a distinct conformation, with the guanine base neither observed to contact residues from the C motif nor the P motif in Box II in their AlphaFold3-predicted model (Supplementary Fig. 7c). In summary, while we uncovered weak tendencies distinguishing ATP from CTP binders, no reliable predictive rule emerged. Predicting nucleotide specificity within the ParB-CTPase fold remains elusive, and a clearer picture will likely require high-resolution experimental structures of binders with their cognate nucleotide ligands.

## Discussion

The discovery that ParB binds and hydrolyses CTP changed our mechanistic understanding of bacterial chromosome segregation and introduced CTP as a regulatory nucleotide alongside ATP and GTP^19^, yet the broader distribution and ligand specificity of the ParB-CTPase fold remained unclear. Here, we show that this fold is widespread across bacteria, archaea, and phages, frequently as orphan homologs encoded on mobile genetic elements. In eukaryotes, in contrast, the most common representative is sulfiredoxin (Srx), which has a well-established enzymatic role in the repair of hyperoxidized peroxiredoxins^53,54^. The near absence of Srx from bacteria, except in cyanobacteria^53^, is also intriguing and likely reflects the reliance on alternative reactive oxygen species defense pathways in prokaryotes. This sharp contrast highlights how a common ancestral fold followed divergent evolutionary paths in prokaryotes vs. eukaryotes.

Our biochemical assays revealed that ParB-CTPase fold-containing proteins can bind CTP, ATP, and, for the first time, GTP, broadening the ligand repertoire of this fold. In this respect, the ParB-CTPase fold resembles other versatile nucleotide-binding folds in biology: the P-loop NTPase fold, which diversified into ATP- and GTP-binding families (but no CTP-binding so far reported) such as helicases, AAA+ ATPases, and Ras proteins^91–94^, and the Rossmann fold, which adapted to bind chemically distinct cofactors such as NAD and SAM^94,95^. The difference is that ParB-CTPase-containing homologs diversify their specificities within the three canonical NTPs themselves. Notably, none of the experimentally tested proteins bound UTP; however, it would not be surprising if ParB-like UTP binders are discovered with deeper future sampling, given the versatility of this fold. Interestingly, 17 of 28 tested proteins showed no detectable NTP binding. This could reflect a genuine loss of nucleotide utilization or a requirement for additional cofactors, such as DNA sequences or partner proteins that enhance nucleotide binding. Moreover, the negative binders might recognize alternative nucleotide derivatives, such as dNTPs, NAD(H), SAM, or cyclic nucleotides. Their persistence across domains of life, however, suggests that they are functional proteins, even if their biochemical activities are not yet fully understood.

Our findings also indicate the ParB-CTPase fold’s capacity to adapt to different cellular roles, by binding different NTP ligands, and raise an important question of whether the ligand choice can be predicted. In canonical ParB, most contacts with CTP involve the phosphate groups, with very few contacts to the cytosine base^5,6,16,17,36,40^. Consistently, AlphaFold3 and molecular docking in our study placed ATP, CTP, and GTP into the same binding pocket, but could not discriminate the correct ligand, likely because current structural prediction tools are trained on existing structural data dominated by CTP-bound ParB, leading to training biases, and because NTPs share identical phosphate-ribose backbones with only differences at the base moiety. Expanding the set of biochemically and structurally validated binders will be crucial for developing predictive rules. These predictive rules, when applied to ParB NTPases across domains of life, would then potentially allow us to answer outstanding questions, such as (i) whether CTP selectivity is ancestral and ATP/GTP preferences evolved later, a scenario requiring fewer evolutionary steps, and whether such switches arose independently multiple times or only once; (ii) which domain architectures are most often combined with ATP- or GTP- or CTP-specific ParB-like folds, and which biological processes they regulate. These insights may eventually explain why and how nucleotide preferences evolved, as well as how they connect to various cellular processes.

Finally, while we identified new ATP-, CTP-, and GTP-binding proteins, binding alone does not prove that these are molecular switches. Canonical ParB and Noc clearly function as CTP-dependent switches^19^, but other homologs, such as SerK, Srx, and PadC, show that the same fold can be used differently, as part of enzymes or as scaffolds, without switch-like conformational changes^6,51,53^. This is not unique to the ParB-CTPase fold but illustrates a broader principle of versatility in protein evolution. The P-loop NTPase fold^93^, for example, spans both ATP- and GTP-binding families and can operate either as a molecular switch (as in GTP-binding proteins Ras^96^ , EF-Tu^97^ , and in ATP-binding proteins ParA^98^, Hsp70 chaperones^99^, or ABC transports^100^) or it can power the translocation of helicases and polymerases through ATP hydrolysis^93^. The protein kinase fold adds another perspective, giving rise to pseudokinases that retain ATP binding but lose catalytic activity, functioning instead as scaffolds in signaling pathways^101^. By analogy, the ParB-CTPase fold may also follow this logic, functioning as a *bona fide* ATP-, GTP-, or CTP-switch in some contexts, or supporting enzymatic functions in others, or persisting as a non-catalytic binder, without any switch activity. In addition to NTPases, many molecular switches rely on cyclic or metabolite-derived ligands such as cAMP, cGMP, c-di-GMP, c-di-AMP, NAD, or SAM, where binding and release, rather than hydrolysis, toggle their activities between on and off state^102–107^. Whether certain ParB-CTPase folds can selectively bind these ubiquitous nucleotides remains to be explored.

Altogether, our findings establish the ParB-CTPase fold as an ancient and versatile nucleotide-binding fold across all life forms, providing a treasure trove of novel proteins whose future characterization may reveal new ways in which living cells and viruses can harness nucleotides to regulate their biology.

## Material and Methods

### Strains, media, and growth conditions

*E. coli* strains were grown in lysogeny broth (LB) medium. When appropriate, the media were supplemented with antibiotics at the following concentrations (μg/mL): carbenicillin (50/100), chloramphenicol (20/30), kanamycin (30/50), streptomycin (50/50), and tetracycline (12.5/12.5).

### Plasmid and strain construction

Strains and plasmids generated or used in this work are listed in Supplementary Tables 2 and 3, respectively. For plasmid construction, a double-stranded DNA fragment containing a desired sequence was chemically synthesized (gBlocks, IDT, Supplementary Table 4). The destination vector was amplified by PCR using oJK45/oJK46 primers (Supplementary Table 4) and purified. A 10 μl reaction mixture was created with 5 μl 2× Gibson master mix (NEB) and 5 μl of a combined equimolar concentration of purified backbone and gBlock(s). Gibson assembly was possible owing to a 20 bp sequence shared between the linearized pTB146 backbone and the gBlocks fragment. This mixture was incubated at 50°C for 60 min. All resulting plasmids were verified by whole-plasmid sequencing (Plasmidsaurus).

### Protein overexpression and purification

Proteins used or generated in this study are listed in Supplementary Table 5.

N-terminally SUMO-tagged ParB-like proteins were expressed from the plasmid pTB146 in *E. coli* Rosetta (BL21 DE3) pLysS competent cells. An overnight culture (10 mL) was used to inoculate 1 L of LB with selective antibiotics. Cultures were incubated at 37 °C with shaking at 220 rpm until the OD_600_ reached ∼0.6. Cultures were cooled for 2 h at 4°C before isopropyl-β-d-1-thiogalactopyranoside (IPTG) was added to a final concentration of 0.5 mM. The cultures were incubated overnight at 16°C with shaking at 220 rpm, after which the cells were pelleted by centrifugation. Cell pellets were resuspended in buffer A (100 mM Tris– HCl, 300 mM NaCl, 10 mM imidazole, 5% (v/v) glycerol, pH 8.0) with 5 mg lysozyme (Merck), 1 μl Benzonase (Merck), and a cOmplete EDTA-free protease inhibitor cocktail tablet (Merck) at room temperature for 30 min with gentle rotation. Cells were lysed on ice with 10 cycles of sonication: 15 s on/15 s off at an amplitude of 20 μm. The lysate was then pelleted at 32,000 g for 35 min at 4°C, and the supernatant was filtered through a 0.22 μm sterile filter (Sartorius). The clear lysate was loaded onto a gravity column containing 2 mL His-Select Cobalt Affinity Gel (Sigma-Aldrich), pre-equilibrated buffer A. Proteins were eluted from the column using 500 mM imidazole in the same buffer. Purified fractions were analysed for purity by SDS-PAGE. The purified protein fractions were loaded onto a preparative-grade HiLoad 10/300 Superdex 200 gel filtration column (GE Healthcare), which had been pre-equilibrated with elution buffer (10 mM Tris–HCl, pH 8.0, 100 mM NaCl, 10 mM MgCl_2_, and 5% (v/v) glycerol). Desired fractions were identified and analysed for purity via sodium dodecyl sulfate–polyacrylamide gel electrophoresis (SDS-PAGE) before being pooled. Aliquots were flash-frozen in liquid nitrogen and stored at -80 °C.

### Differential Radial Capillary Action of Ligand Assay (DRaCALA)

Purified 6xhis-SUMO ParB-like proteins (WT and mutants, at final concentrations of 25 μM) were incubated with 20 μCi/mL of radiolabeled ^32^P-α-NTP (Perkin Elmer), 35 μM of unlabeled NTP (ThermoFisher), in the binding buffer (100 mM Tris pH 8.0, 100 mM NaCl, 10 mM MgCl_2_, 1 mM CaCl_2,_ and 0.5 µM of β-mercaptoethanol) for 10 min at room temperature. Then, 4 μL of the samples were spotted onto a nitrocellulose membrane and air-dried till the spots turned completely invisible on the membrane. The nitrocellulose membrane was put in plastic pouches and then exposed to a phosphor screen (Fujifilm) for 10 minutes. Each DRaCALA assay was duplicated, and a representative autoradiograph is shown.

### AlphaFold 3 structural predictions and molecular docking of NTPs

AlphaFold 3 was used to predict the structures of ParB-like protein complexes with various NTPs^108^ . Briefly, the input files were populated with the sequences of WT ParB-like proteins and NTP structures supplied via Chemical Component Dictionary codes. The program was run with default parameters, with both data and inference pipeline enabled. Then, to obtain a more accurate description of NTP’s position and an estimate of the binding energy, AutoDock GPU was used (^109^ and <u>GitHub - ccsb-scripps/AutoDock-GPU:</u> <u>AutoDock for GPUs and other accelerators</u>). Firstly, to prepare for the docking process, the AlphaFold 3 models were processed with PROPKA3 to assign the charge and protonation state of side chains corresponding to pH 7.4^110^. Then, the NTPs were docked into the protonated molecule using the AutoDock 4 force field, and the best prediction was presented.

### Homology survey of ParB and ParB-CTPase fold-containing proteins and associated PFAM domains

Phage proteomes were obtained from the Gut Phage database^111^, and the Millard laboratory repository^112^. A local bacterial and archaeal genome database was built following the GTDB taxonomic framework (version 220). The version used in this work is an updated version from Strock et al. 2025^55^. Eukaryotic proteomes were obtained from EukProt v3. TIGR, PFAM, and KOFAM HMM models of ParB homologues, ParB-CTPase fold, and ParA (TIGR00180.1, PF02195.22, and K03496, respectively) were searched against each proteome database using hmmsearch (HMMER v3.4, option --cut_ga except for K03496, e-value <1e-3). All other PFAM domain hits were computed using the hmmsearch option cut-ga, against the PFAM-A database (downloaded on 23.12.01)

### *parAB* operons and mobile element detection

For each species, prophage and plasmids were predicted using GeNomad^113^ (end-to-end mode, Docker image obtained on 25.04.30). We annotated a ParB protein as part of a ParAB pair when its encoding gene was located two genes downstream or upstream from a *parA* homolog.

### Alignments and phylogenetic trees

All species trees were obtained from the GTDB. Both the ParB homolog and ParB-CTPase domain tree were built on a reduced-size database, restricted to one species per family. For the ParB-CTP-binding domain tree, only prophages from the Millard Laboratory database were considered. Proteins were aligned using MAFFT v7.490, and alignments clipped using Clipkit v2.25 (mode gappy). ParB homologs -TIGR00180- tree was built using IQ-TREE multicore version 2.2.5 (-alrt 1000 -nt AUTO -mem 32G -m TEST -s). Owing to size considerations, the ParB-CTPase fold tree was built using fastree v2.1.11. ParB-CTPase fold tree initial sequence set was obtained from the alignment output of hmmsearch, which was subsequently re-aligned and trimmed. For all the alignments the trimming was done using default setting (from the manual: (default: 0.9).

### Sequence co-evolution and ParB-CTPase fold motifs

All sequence logos were plotted using R, ggseqlogo, excluding alignment gaps. To compute the residues most associated with overall divergence, we first applied a principal component analysis to a distance matrix computed from the protein alignment. Next, we computed the squared correlation between each amino acid (binary encoded) and the first principal component of the alignment. For each position, we computed the sum of all squared correlations as a contribution score.

## Data availability

All protein sequences, alignments, and phylogenetic trees are available on Zenodo (https://zenodo.org/records/17224332). Scripts used for this study are available on GitHub (https://github.com/HocherLab/ParB_Evo_Github).

## Acknowledgement

We thank Romain Strock for kindly sharing an updated version of the prokaryotic proteome database. We are grateful to all members of the Tung Le lab, Mark J Buttner, Susan Schlimpert, Max Jordan, Martin Thanbichler, and Geraldine Laloux for insightful discussions and thorough reading of the manuscript. This work is supported by a Lister Institute Fellowship and Wellcome Trust Investigator grant 221776/Z/2/Z (to T.B.K.L.) that funds J.K., K.V. S., and K. E. J., and the BBSRC-funded Harnessing Biosynthesis for Sustainable Food and Health (HBio) Institute Strategic Programme BB/X01097X/1 (to the John Innes Centre). A.H is supported by a Wellcome Trust Career Development Award (227755/Z/23/Z).

## Authors contributions

Conceptualization: JK, AH, TBKL

Methodology: JK, KVS, AH

Investigation: JK, KVS, KEJ, AH, TBKL

Visualization: JK, KVS, AH

Funding acquisition: AH, TBKL

Project administration: JK, TBKL

Supervision: TBKL

Writing - original draft: JK, AH

Writing - review & editing: JK, KVS, AH, TBKL

---

**Supplementary Table 1.** Distribution of chromosomal *parB* across bacterial orders with near-complete genomes. Only orders in which >90% of species encoded chromosomal *parB* are shown. The table lists, for each order, the number of species with or without chromosomal *parB*, along with the corresponding percentage.

**Supplementary Table 2.** Description of bacterial strains used in this study.

**Supplementary Table 3.** Description of plasmid constructions used in this study.

**Supplementary Table 4.** Description of oligonucleotides used in this study.

**Supplementary Table 5.** Description of proteins purified and analyzed in this study.

**Supplementary Table 6.** Details of Box I (C motif) and Box II (P motif) and predicted nucleotide contact residues in ParB-NTP proteins based on molecular docking.

**Supplementary Fig. 1.**
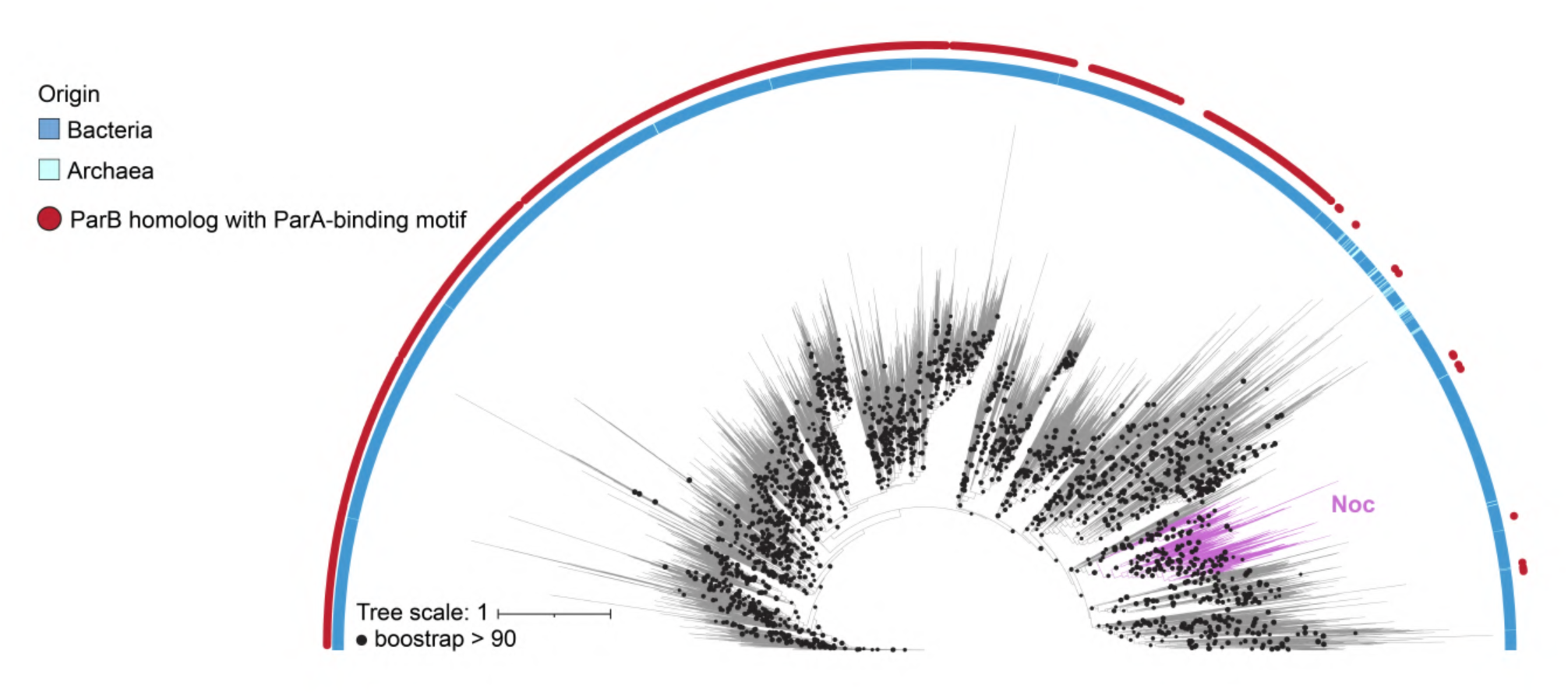
Distribution of ParB proteins across bacteria and archaea. Related to. **Fig. 2-3**. Midpoint rooted single gene phylogenetic bee of bacterial and archaeal ParB homologs (TIGR00180, see Methods). The inner ring indicates the taxonomic origin: blue represents bacteria, and yan represents archaea. The outer red ring marks ParB homologues with the ParA-binding motif. Black dots denote nodes with ultrafast bootstrap support ;?90. The Noc protein, an example of an orphan ParB homolog that lost the ParA-binding motif, is indicated in magenta.

**Supplementary Fig. 2.**
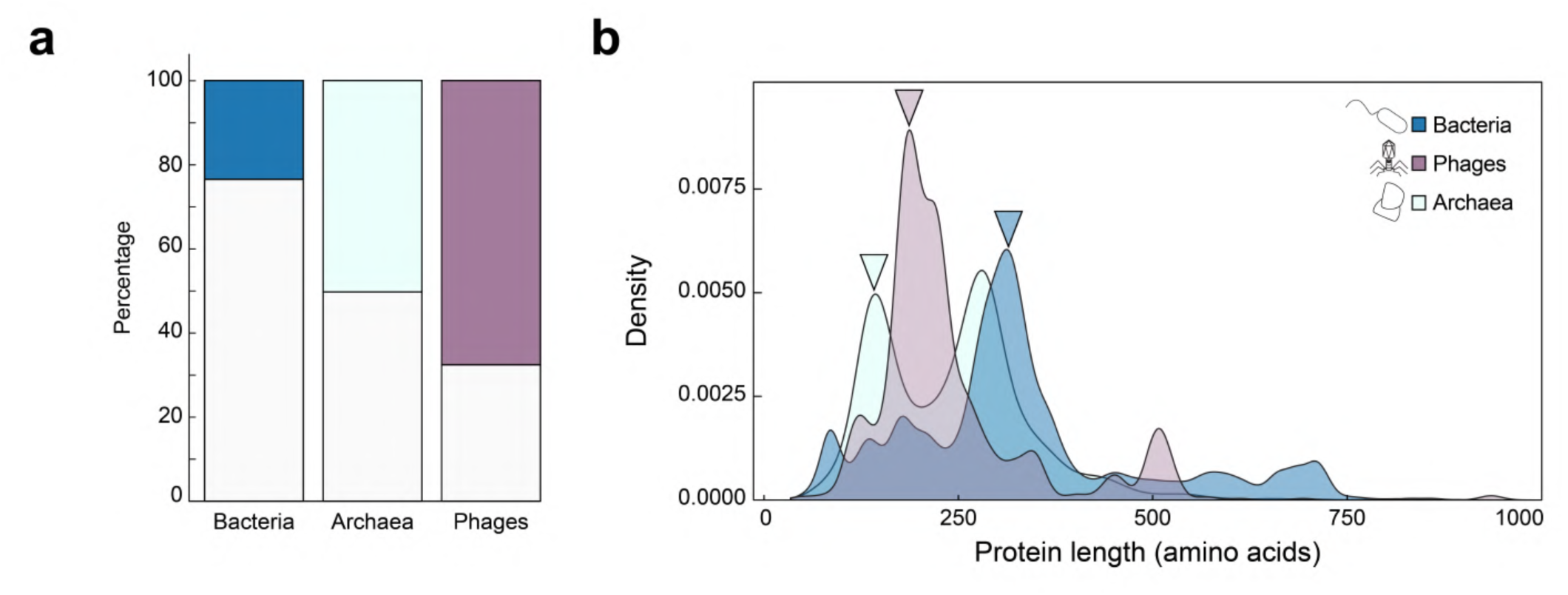
Length distribution of proteins that are annotated as single-domain ParB-CTPase fold-containing homologs. Related to. **Fig. 4. a** Fraction of ParB-CTPase fold-containing proteins annotated as “single-domain” (colored bars) versus multi-domain (grey bars, as shown in Fig. 4a) in bacteria, archaea, and phages. **b** Protein length distribution of the “single-domain” proteins subset, shown as density plots for bacteria (blue), archaea (cyan), and phages (purple). Short proteins (−100-200 amino acids) dominate in phages and some archaea (magenta and cyan arrowheads), consistent with a single ParB-CTPase fold. In contrast, many bacterial proteins exceed 300 amino acids {blue arrowhead), which is longer than the ParB-CTPase fold itself, suggesting that they likely contain additional domains that were not annotated.

**Supplementary Fig. 3.**
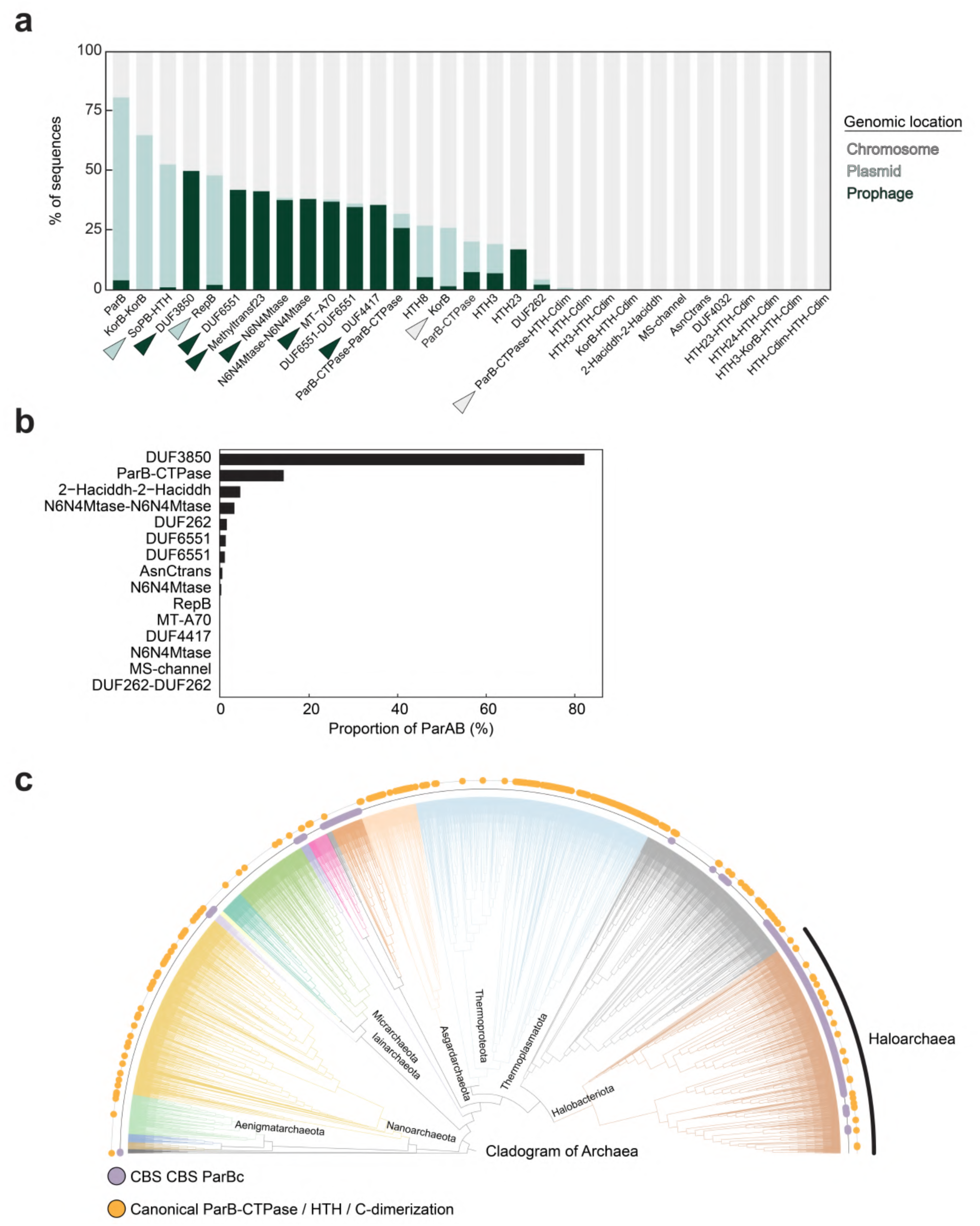
Distribution and genomic context of genes encoding for novel ParB-CTPase fold-containing proteins. Related to. **Fig. 4. a** Genomic location of ParB-CTPase fold-containing proteins. The x-axis lists ParB-CTPase fold-containing proteins fused to different domains. The y-axis shows the percentage of sequences for each fusion type encoded on chromosomes (grey), plasmids (blue), or prophages (dark green). Examples highlighted in text are marked with arrowheads. **b** Association of ParB-CTPase fold-containing proteins with the *parA* gene. Bars show the percentage of proteins that are encoded adjacent to a *parA* homolog for each domain fusion. ParB-CTPase-DUF3850 proteins are most frequently found in operons with *parA.* c Cladogram of archaeal species showing species encoding ParB-CTPase fold fused with CBS-domain (purple dots in the inner ring) and encoding ParB-CTPase fold fused with canonical DNA binding motif HTH and C-terminal dimerisation domains (orange dots in outer ring). CBS fusions are confined to specific clades, whereas canonical ParB proteins occur more patchily and typically are not encoded with a *parA* gene.

**Supplementary Fig. 4.**
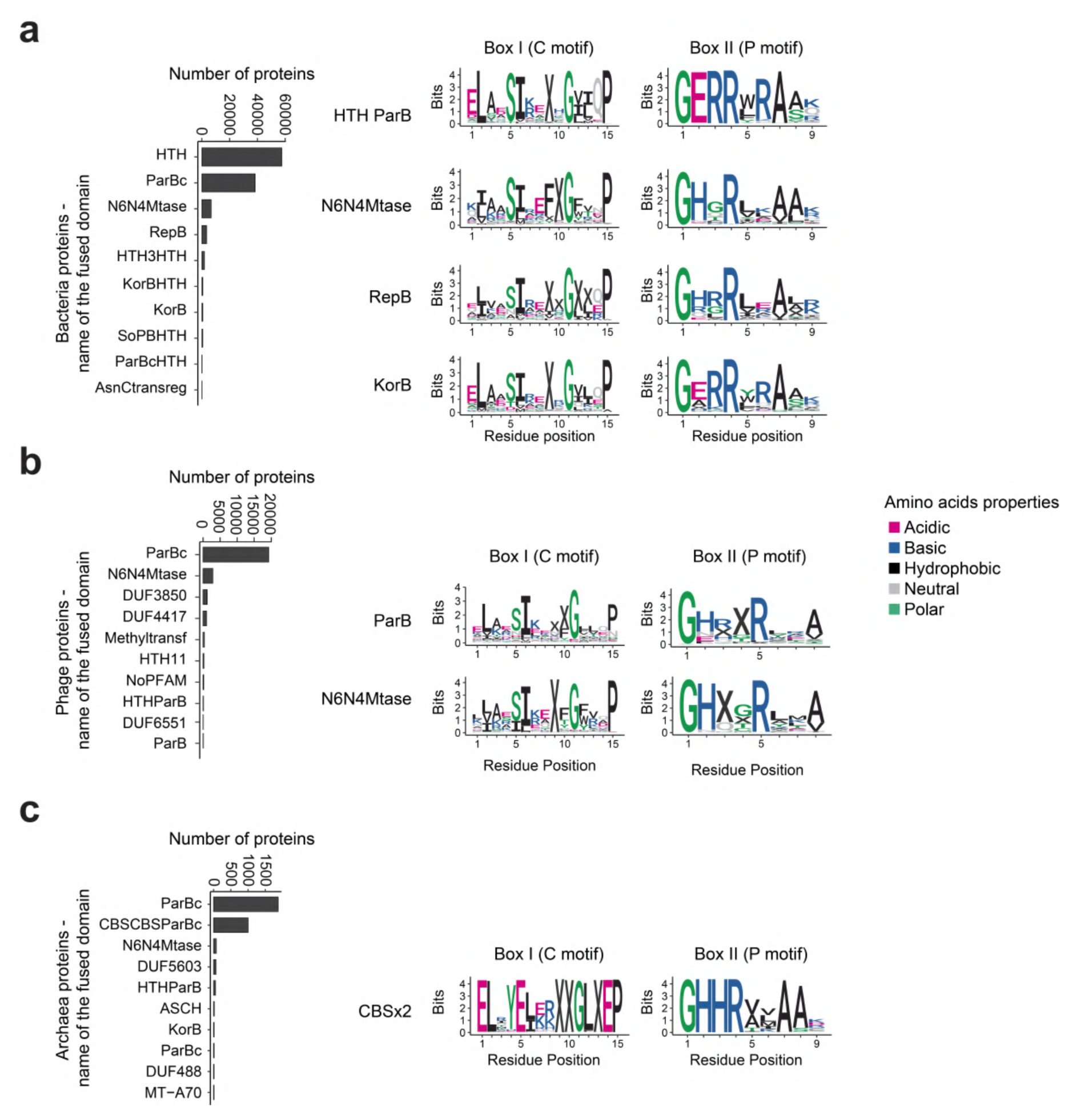
Subtle variation of ParB-CTPase motifs across different ParB-CTPase fold-containing proteins. Related to. **Fig. 4. a** Bacterial proteins. Left: frequency of domains fused to the ParB-CTPase fold. Each bar represents the number of proteins carrying the indicated fusion domain (higher bars = more frequent fusion). Right: sequence logos showing conserved Box I (C motif) and Box II (P motif) motifs for selected, most common families (e.g., HTH, N6N4-methyltransferase, RepB, KorB). b Phage proteins. Left: most frequent ParB-CTPase fusions in phages. Right: Box I (C motif) and Box II (P motif) logos for common families such as stand-alone ParB-CTPase fold proteins and N6N4-methyltransferases. cArchaeal proteins. Left: most frequent ParB-CTPase fusions in archaea. Right: Box I (C motif) and Box II (P motif) logos for the most common CBS domain fusion. Amino acids are colored based on their chemical properties.

**Supplementary Fig. 5.**
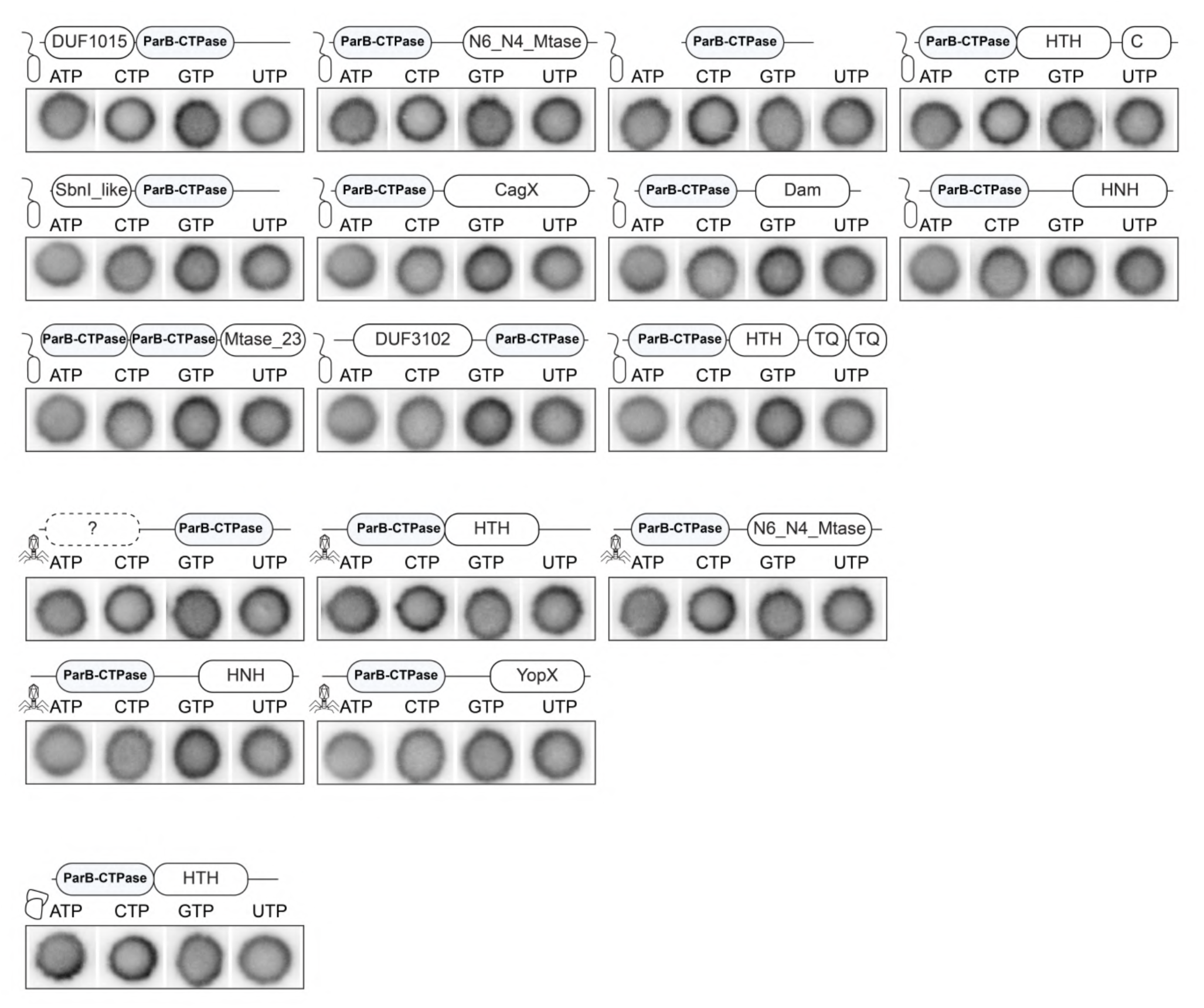
Screening ParB-CTPase fold-containing proteins for nucleotide binding using DRaCALA. Related to. **Figure 5**. Different ParB-CTPase fold-containingproteins were tested for binding to radiolabeled NTP a-P32, and none showed NTP-binding. The starting concentration of proteins used was 25 µM. DRaCALA assays were performed with two replicates; one representative replicate is shown. Cartoon icons indicate the origin of each protein (phage, bacterium, or archaea). Domain schematics above each protein indicate representative fusions, with the ParB-CTPase domain shown in blue and fused to various domains. Proteins are grouped by their origin. Detailed information for each protein (including source organism, accession numbers, and domain annotations) is provided in Supplementary Table 5.

**Supplementary Fig. 6.**
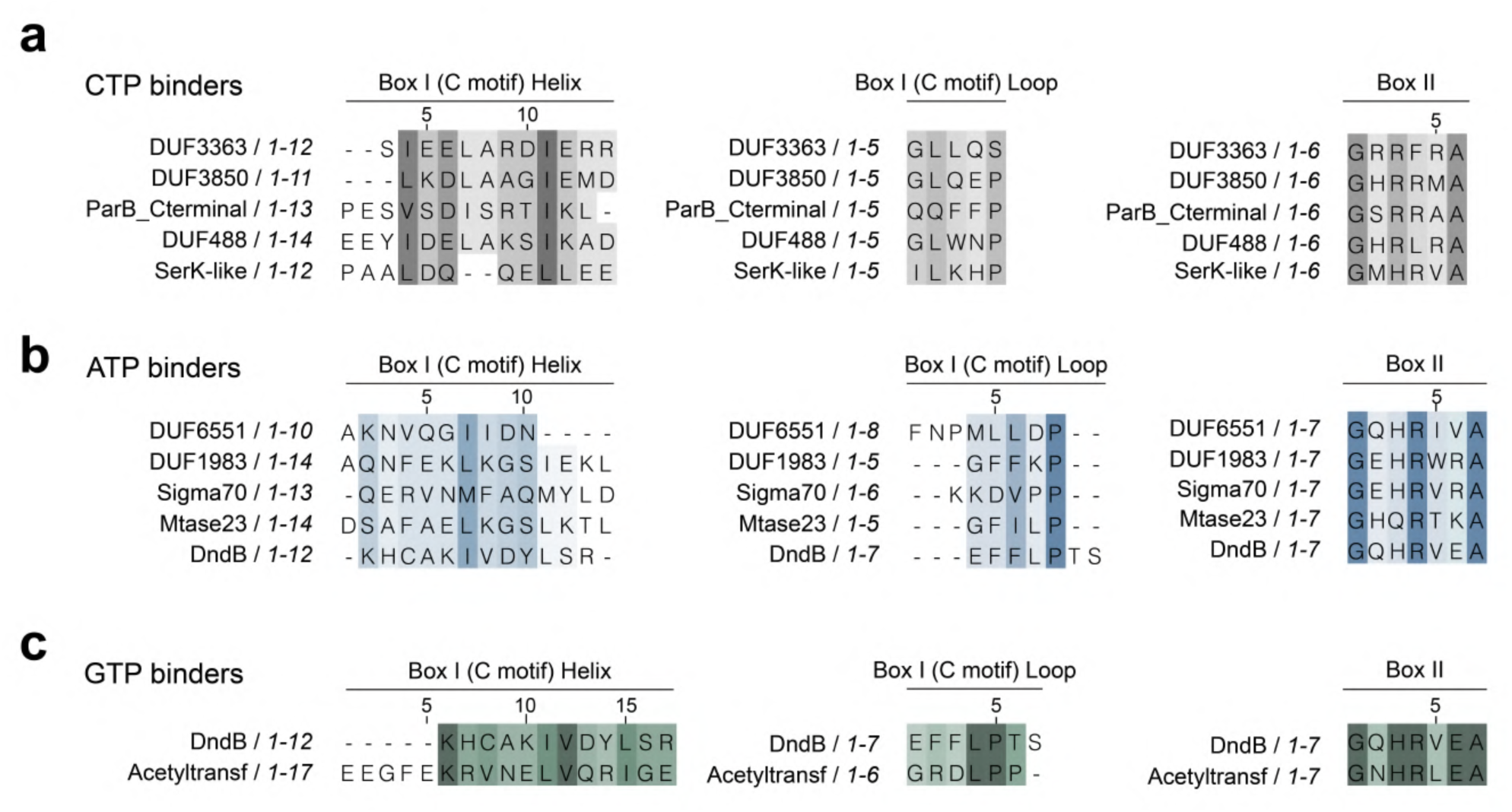
Sequence alignments of conserved motifs in ATP-, CTP-, and GTP-binding ParB-like proteins. Related to Fig. 6. For each nucleotide-binding protein, AlphaFold2 structural models were used to identify Box I Helix, Box I Loop (C motif), and Box II (P motif). The sequences of these motifs were then extracted and aligned separately for CTP-, ATP-, and GTP-binding proteins. Alignments were performed using MUSCLE and visualized in Jalview, with residues coloured by conservation. **a** CTP binders (grey). Representative proteins include ParB-CTPase fold fused with DUF3363, DUF3850, ParB_C-terminal, DUF488, and SerK-like domains. **b** ATP binders (blue). Representative proteins include ParB-CTPase fold fused with DUF6551, DUF1983, Sigma70, Mtase23, and DndB domains. **c** GTP binders (green). Representative proteins include ParB-CTPase fold fused with DndB and Acetyltransferase domains. These alignments highlight subtle sequence features across binding classes and were used in preparation for Figs. 6a and 6b. See also Supplementary Table 6.

**Supplementary Fig. 7.**
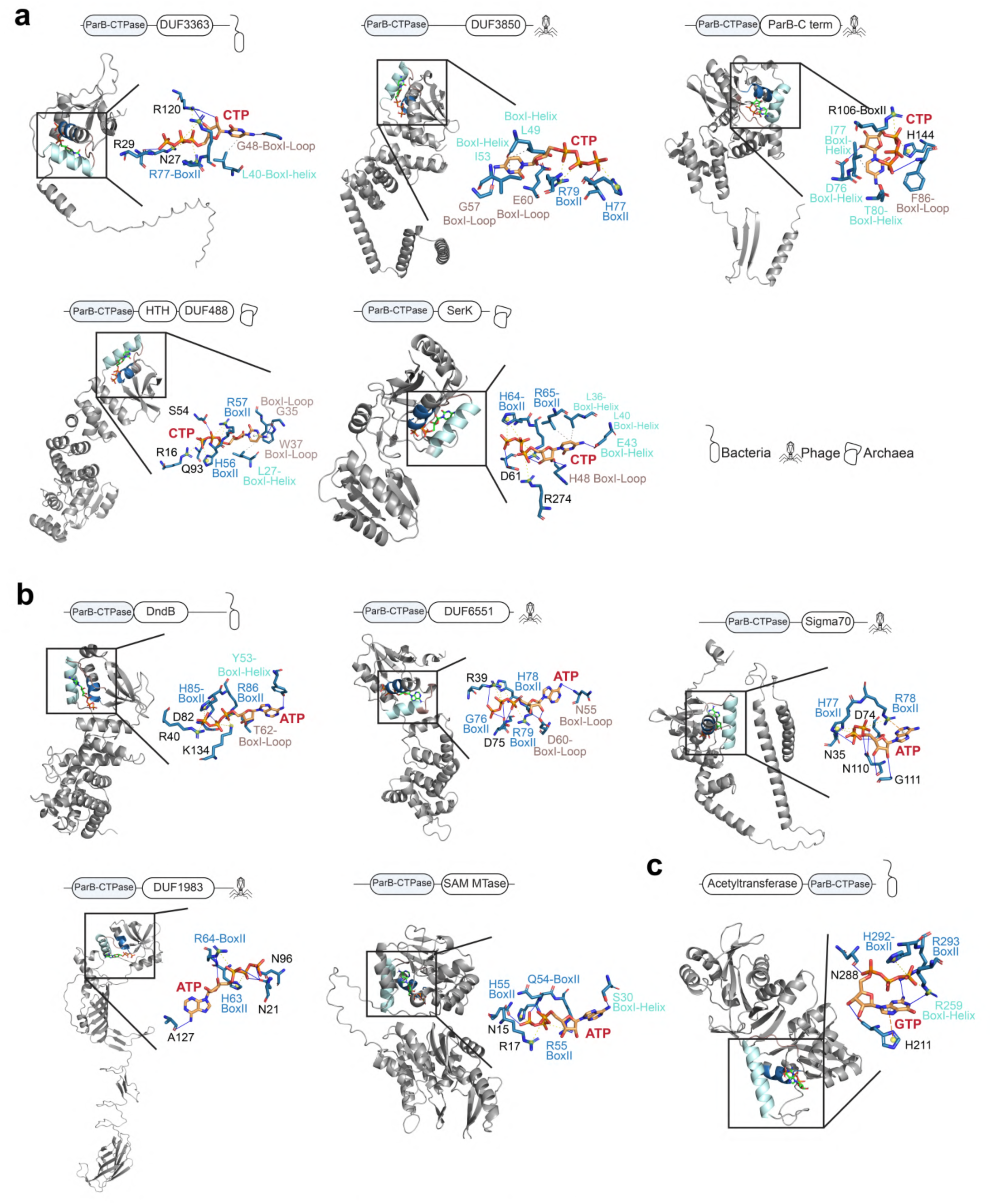
Related to Fig. 6. Structural predictions and docking of ATP-, CTP-, and GTP-binding ParB-CTPase fold-containing proteins. **a-c** AlphaFold2 models of the 11 ParB-CTPase fold proteins that showed positive nucleotide binding in DRaCALA were used for molecular docking with their respective ligands. Proteins are grouped by binding preference: **a** CTP binders, **b** ATP binders, **c** GTP binders. For each protein, the schematics on the top show domain organization, with the ParB-CTPase fold in blue and fused domains in white. Icons indicate the source of the protein (bacteria, phage, or archaea). In the structural models, the ParB-CTPase fold is shown with Box I Helix (C motif) in cyan, Box I Loop (C motif) in maroon, and Box II (P motif) in blue. Insets show zoomed docking poses of the ligand (CTP, ATP, or GTP, in red) and predicted interacting residues. Residues are labelled according to whether they originate from Box I Helix and Box I Loop (C motif), or Box II (P motif). See also Supplementary Table 6.

**Supplementary Fig. 8.**
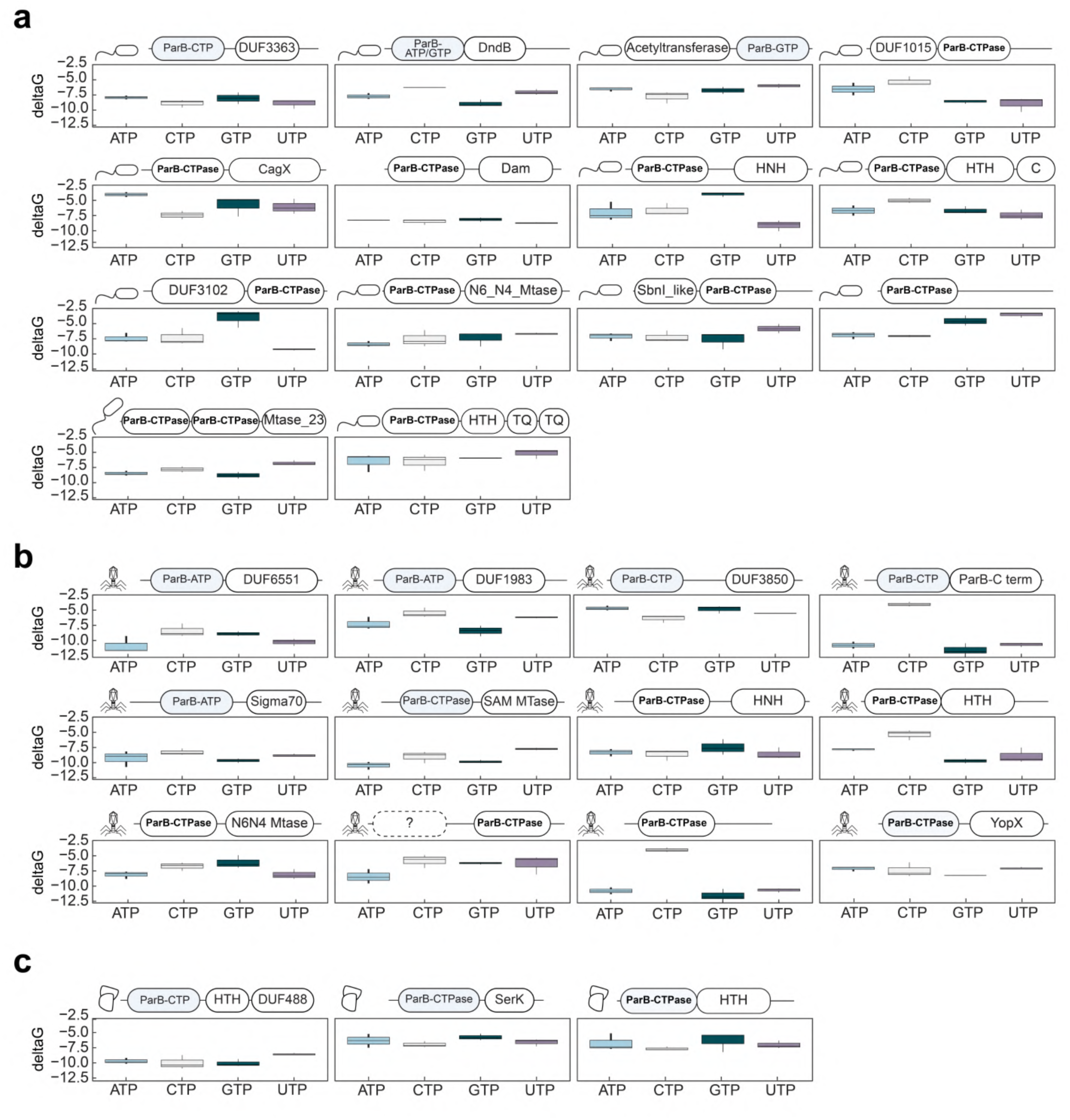
AlphaFold3-predicted docking energies of nucleotides to ParB-CTPase fold-containing proteins. Related to Fig. 6. Predicted binding energies (ΔG. kcal/mol) for ATP, CTP, GTP, and UTP were obtained using AlphaFold3 docking for all proteins tested in DRaCALA assays (positive and negative binders; see Fig. 5 and Supplementary Fig. 5). Domain architectures are shown on top of the corresponding docking scores. For positive binders, the ParB-CTPase fold is highlighted in blue, and the experimentally confirmed ligand is written. For negative binders, the ParB-CTPase fold is shown in white. Proteins are grouped by origin: bacteria (a), archaea (b), and phages (c).

